# Targeted insertion and reporter transgene activity at a gene safe harbor of the human blood fluke, *Schistosoma mansoni*

**DOI:** 10.1101/2022.09.02.506379

**Authors:** Wannaporn Ittiprasert, Max F. Moescheid, Cristian Chaparro, Victoria H. Mann, Thomas Quack, Rutchanee Rodpai, André Miller, Prapakorn Wistiphongpun, Watunyoo Buakaew, Margaret Mentink-Kane, Sarah Schmid, Anastas Popratiloff, Christoph G. Grevelding, Christoph Grunau, Paul J. Brindley

**Affiliations:** Department of Microbiology, Immunology & Tropical Medicine, School of Medicine & Health Sciences, George Washington University, Washington, D.C. 20037; Institute of Parasitology, Biomedical Research Center Seltersberg, Justus Liebig University Giessen, Giessen, Germany; IHPE, Univ Perpignan Via Domitia, CNRS, IFREMER, Univ Montpellier, Perpignan, France; Department of Parasitology and Excellence in Medical Innovation, and Technology Research Group, Faculty of Medicine, Khon Kaen University, Khon Kaen 40002, Thailand; Schistosomiasis Resource Center, Biomedical Research Institute, Rockville, MD, 20850; Faculty of Medical Technology, Rangsit University, Pathum Thani, 12000, Thailand; Department of Microbiology, Faculty of Medicine, Srinakharinwirot University, Bangkok, 10110 Thailand; Nanofabrication and Imaging Center, Science & Engineering Hall, George Washington University, Washington, D.C. 20052

**Keywords:** overlapping CRISPR target, reporter transgene, gene safe harbor, human blood fluke, *Schistosoma mansoni*

## Abstract

The identification and characterization of genomic safe harbor sites (GSH) aims to facilitate consistent transgene activity without disruption to the host cell genome. We combined genome annotation and chromatin structure analysis by computational approach to predict the location of four GSHs in the human blood fluke, *Schistosoma mansoni*, a major infectious pathogen of the tropics. Introduction of a transgene into the egg of the parasite was accomplished using CRISPR/Cas-assisted homology-directed repair and overlapping guide RNAs. Gene editing efficiencies of 24% and transgene-encoded fluorescence of 75% of gene-edited schistosome eggs were observed. These outcomes advance functional genomics for schistosomes by providing a tractable path towards transgenic helminths using homology directed repair-catalyzed transgene insertion. This approach should be adaptable to helminths generally.

**Motivation:** Functional genomics methods are needed in the arena of the Neglected Tropical Diseases, especially for helminth parasites, to facilitate basic and transformational studies to improve global public health. Gain-of-function phenotypes would be valuable in this context. Hence, the motivation for this investigation was to identify a genome safe harbor (GSH) site in the chromosomes of the human blood fluke, *Schistosoma mansoni* and, in a pilot approach, and to develop methods for transgene insertion and methods to characterize transgene performance following homology directed insertion at the schistosome GSH. The progress reported here can be expected to advance functional genomics for species of the Platyhelminthes – including model, free living species of planarians, and should be adaptable to helminths generally.

## Introduction

Clustered Regularly Interspaced Short Palindromic Repeats (CRISPR) technology has revolutionized functional genomics in biology, medicine and agriculture ^1–3^. Transgenesis approaches are integral in diverse applications including gene therapy, biotherapeutics and deciphering host-pathogen interactions. With progress emanating from model species and cell lines, tools and techniques can frequently be adapted and transferred to non-model species. Among these are the parasitic helminths responsible for several major neglected tropical diseases (NTDs), which cause substantial morbidity and mortality. NTDs mainly occur in the Global South, and they are responsible for a disease burden that exceeds that caused by malaria and tuberculosis ^4^. Infections with parasitic helminths also are responsible for substantial economic and disease burdens in the agriculture and animal health ^5^. These public health and economic imperatives motivated international collaboration for parasite *omics* research that has produced outsized databases of genomes and proteomes, and gene, transcript, and protein annotations ^6–9^. In the current post-genomics era, and despite the availability of these omics data, tools for functional genomics in parasitic helminths have been limited to RNA interference, which performs with variable efficacy^10, 11^. Therefore, access to CRISPR-based transgenesis protocols for functionally characterizing genes of interest such as those coding for putative drug and/or vaccine targets has eminent research priority. Moreover, progress with gene editing technology in schistosomes will facilitate its use in other major invertebrate clades of the Protostomia, including the planarians, for which CRISPR-based reverse and forward genetics have yet to be reported.

CRISPR enables targeted site-specific mutation(s), obviating an impediment of earlier transgenesis approaches that relied on vector-based particle bombardment^12^, lentiviruses^13^ and transposons such as *piggyBac*^14^. These latter approaches could lead to genetic instability, multi-copy insertion, unstable expression or inactivation of the transgene and interference with the endogenous gene under investigation. These issues can be overcome in the process of genome editing, where double stranded breaks (DSBs) are resolved by several discrete repair mechanisms, particularly the predominant error-prone non-homology end joining (NHEJ) and the templated homologous-directed repair (HDR) pathways. Sister chromatids provide a natural repair template, whereas exogenous DNA such a plasmid, oligodeoxynucleotides, and PCR amplicons can also serve as the repair template. HDR efficiency can be markedly improved when supplied with double-strand (ds) donor DNA with modifications^15^. CRISPR/Cas-assisted HDR has been applied in *Schistosoma mansoni*^16, 17^ with promoter-free, single strand-deoxynucleotide donors. Multiple overlapping CRISPR target sites improve precise HDR insertion of large cargoes in embryonic stem cells^18, 19^, while modification of 5’-termini of long dsDNA donors enhance HDR, bolstering efficient, single-copy integration through the retention of a monomeric donor confirmation and thereby enabling gene replacement and tagging^20^.

One of the caveats of transgene integration is that their insertion into arbitrarily chosen positions in the genome may lead to loss of expression due to disruption of cell function or repressive chromatin structure in the target region. This had been identified as a major drawback, initially in gene therapy approaches, and has led to the concept of Genome Safe Harbors (GSHs)^21–24^. An ideal GSH has been defined as a region (i) that does not overlap (predicted) functional DNA elements and (ii) without heterochromatic marks that could impede transcription^25^. This approach was successfully used in model organisms such as *Caenorhabditis elegans* based on annotations from the ENCODE and modENCODE consortia^26^. For non-model organisms, chromatin structure annotations are often not available and experimenters resort to criterion (i). For instance, transgene insertion into a GSH of the human filarial parasite *Brugia malayi* has been reported but, in this case, GSHs were predicted based on four sequence annotation features alone: located in intergenic regions, to be unique in the genome, to contain a terminal protospacer adjacent motif (PAM) necessary for recognition by the sgRNA/CAS9 RNP, and fourth, the putative PAM sequence admissible only if situated > 2 kb from the nearest predicted coding region^27^.

In this study, we profited from the availability of chromatin data (ChIP-Seq of histone modifications), chromatin accessibility data (ATAC-Seq and HIV integration) and combined them by a computational investigation with genome sequence information to identify potential GSH sites in *S. mansoni*. Furthermore, we adapted CRISPR/Cas9-based approaches to insert a reporter transgene into the most qualified out of four predicted candidate GSHs, which is situated in an euchromatic region of chromosome 3. The donor transgene encoded enhanced green fluorescent protein (EGFP) under the control of the schistosome ubiquitin gene promoter and terminator. The targeted region was free of repetitive sequences and neighboring long non-coding regions, a situation likely to minimize off-target effects of CRISPR/Cas activity. Multiple PAMs within this region were targeted with overlapping single guide RNAs, deployed in unison to enhance editing efficiency and homology directed repair (HDR) in the presence of the phosphorothioate-modified, reporter transgene donor DNA template. A knock-in (KI) efficiency of 75% was observed by reporter transgene-mediated expression of EGFP in miracidia developing within the schistosome eggshell.

## Results

### Genome safe harbors predicted in the schistosome genome

To identify potential GSH sites we performed a combination of *in silico* analyses based on accepted criteria, newly introduced principles ^28^, and genome sequence resources available for *Schistosoma mansoni*, which could satisfy benign and stable gene expression. Notably, we sought to identify intergenic GSH, rather than intragenic GSH ^28^. Four regions satisfied our criteria (Fig. 1A; detailed in the Methods) and were termed GSH1 (1,416 bp in length; location, chromosome 3:13380432-13381848), GSH2 (970 bp; chromosome 2: 15434976-15435954), GSH3 (752 bp; chromosome 2: 9689988-9690739), and GSH4 (138 bp; chromosome 3: 13381901-13382038), respectively (Fig. 1), the naming based on the rank order of their sizes from longest (GSH1) to the shortest GSH (GSH4). We note that several protein-coding gene loci were situated in the vicinity of these gene-free GSHs, although these genes were > 2kb distant from any GSH: Smp_052890, uncharacterized protein; Smp_150460, copper transport protein; Smp_071830, uncharacterized protein; Smp_245610, uncharacterized protein and Smp_131070, condensing complex (Figs. 1B-D). Most or all of these genes are as yet uncharacterized proteins, and may be non-essential genes based on orthology to essential genes known from model eukaryotes ^29^. For CRISPR-specific considerations for the programmed transgene insertion, particularly the presence of multiple PAMs, GSH1 qualified as likely the most useful of the four GSH for the present investigation and hence programmed gene editing at GSH1 is the focus of the findings detailed below.

**Figure. 1.**
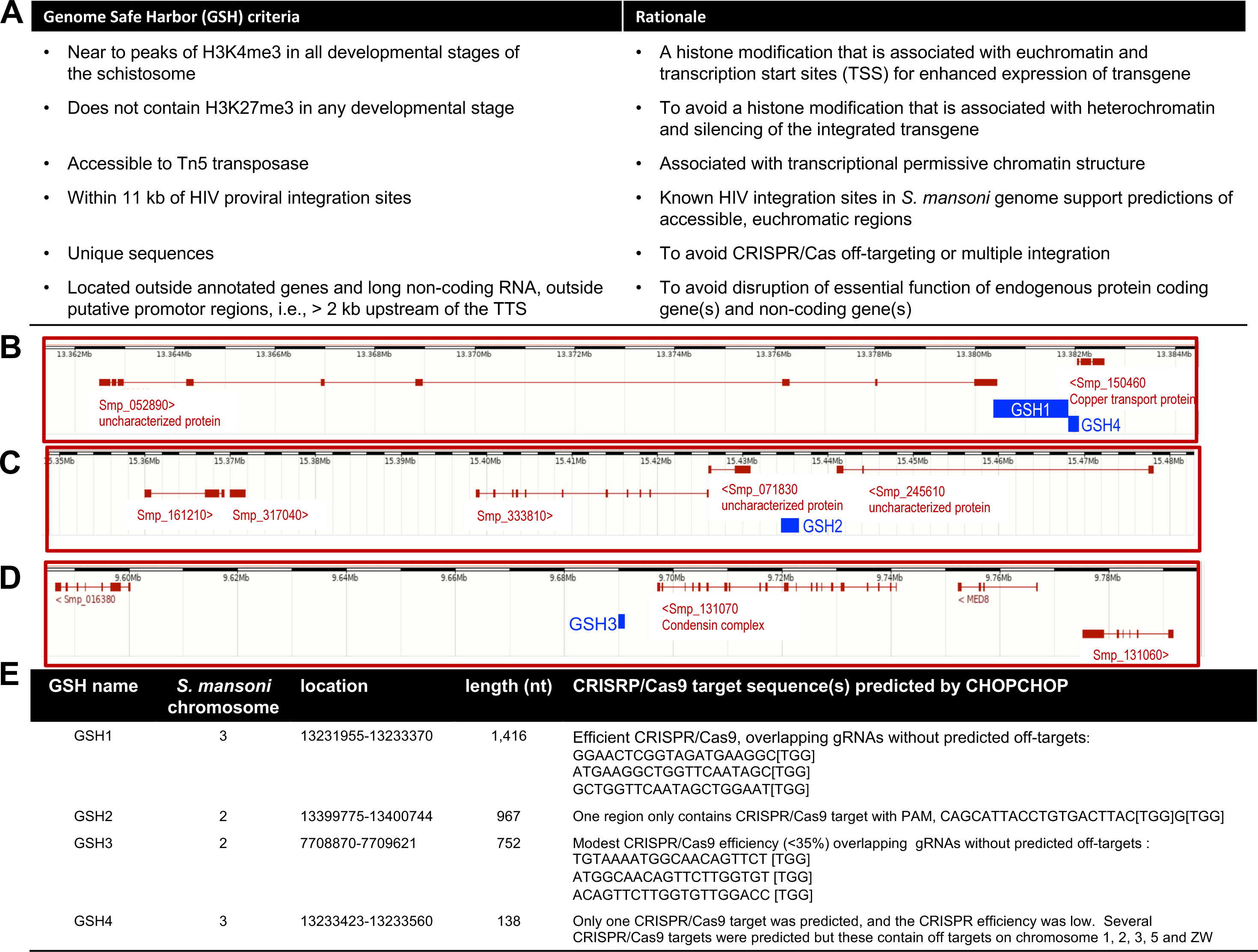
Bioinformatic identification of GSHs and their CRISPR-programmed targets in the genome of S. mansoni. Panel A, Criteria and rationale used to computationally predict genome safe harbors (GSHs). Panels B-D, Chromosomal locations and lengths of four candidate GSH sites (blue boxes), located on chromosome 2 (GSH2 and GSH3) and chromosome 3 (GSH1, GSH4) of *S. mansoni* (annotated using *S. mansoni* genome v9). All criteria were equally weighted. In addition, coding sequences of genes identified in the vicinity of the four (gene-free) GSHs are shown, which were mostly uncharacterized proteins (red boxes with connected red line). Black and white scale bars at the top of each panel each represent one megabase pairs in length. Panel E, Features of the GSHs including chromosomal location, length, programmed CRISPR/Cas9 targets, guide RNAs, and their predicted on- and off-target specificities from the CHOPCHOP design toolbox.

### Efficiency of programmed mutation at GSH1 enhanced by multiple guide RNAs

Having predicted the locations of these GSHs, we proceeded to investigate the efficiency of programmed mutation and reporter transgene activity at GSH1, chromosome 3: 13380432-13381848. Overlapping guide RNAs were employed, an approach that enhanced KI efficiency in some mammalian cell lines and embryos ^18, 19^. Among the guide RNAs (gRNA) exhibiting on-target specificity for GSH1, three overlapping synthetic guide RNAs; sgRNA1, sgRNA2 and sgRNA3, which lacked self-complementarity and off-target matches to the reference *S. mansoni* genome (Figs. 1B, E, Fig. 2A), were selected from among CRISPR/Cas9 target sites predicted by CHOPCHOP ^30, 31^. The ribonucleoprotein complexes (RNPs) of Cas9 nuclease and sgRNA were assembled, after which four discrete mixtures of the three RNPs were used. Three of the mixtures included two RNPs, *i.e.* dual RNPs mixtures (RNP1+RNP2, RNP2+RNP3, and RNP1+RNP3) and the fourth included the three RNPs, *i.e.* triple RNPs (RNP1+RNP2+RNP3).

**Figure. 2.**
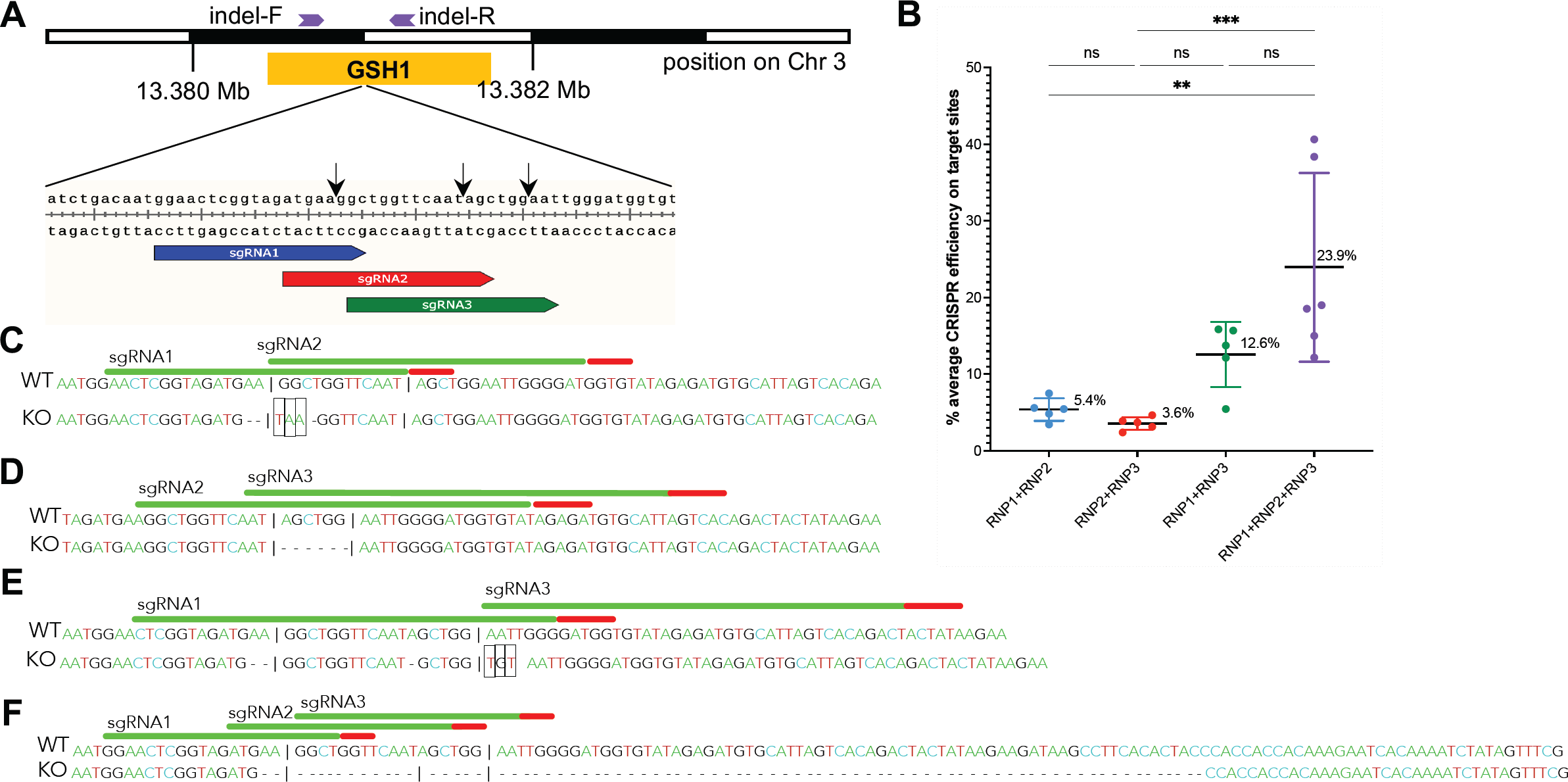
Efficiency of programmed knockout enhanced using three overlapping guide RNAs. Panel A, Schematic map of sites of the overlapping guide RNAs (blue, red and green arrows for target 1, -2 and -3, respectively) within GSH1 (yellow box), along with primer locations for indel analysis (purple arrows). The black arrows indicate the site programmed double stranded break (DSB) programmed by sgRNA1, 2 and 3. Panel B, Efficiency of CRISPR knockout at GSH1 in eggs of *S. mansoni*, as assessed with the Deconvolution of Complex DNA Repair (DECODR) algorithm using distance, following transfection with overlapping guide RNPs; RNP1+RNP2 (blue dots), RNP2+RNP3 (red), RNP1+RNP3 (green), and RNP1+RNP2+RNP3 (purple). Significantly higher CRISPR efficiency obtained with the three-overlapping guide RNPs, mean = 23.6%, than the other groups (*P* ≤ 0.001). Among the groups transfected with dual RNPs, efficiency obtained with the RNP1+RNP3 treatment group, mean = 12.6%, was significantly higher than either of the other groups, RNP1+RNP2 at 5.4% and RNP2+RNP3 at 3.6% (*P* ≤ 0.01; one-way ANOVA with 95% confidence intervals, five or six independent biological replicates; GraphPad Prism). Panels C-E, Representative alleles, in the schistosome genome, bearing indels at the target site in GSH1 (panel A) following transfection with dual guide RNPs, as a gauge of efficiency CRISPR-catalyzed gene editing. The reference wild type (WT) allele is shown above the mutant (KO) allele. KO alleles were identified during examination of nucleotide sequences for small deletions, 1-6 nt in length (-dash) or insertion/substitution (black boxes) The vertical black line boxes show targeted PAM (protospacer adjacent motif) trinucleotides. Panel F, A representative example of an KO allele bearing a large sized deletion resulting from transfection with the triple guide RNPs.

The mixtures of these multiple RNPs, along with the donor DNA template encoding EGFP, was delivered to schistosome eggs by electroporation. The transfected eggs were maintained in culture for 15 days, after which EGFP expression was quantified. Thereafter, genomic DNAs and total RNAs were extracted from the eggs. Efficiency of genome editing, both in mock-treated controls and experimental groups, was assessed using DECODR ^32^ analysis of Sanger sequence chromatograms of amplicons that spanned the CRISPR/Cas9 programmed DSBs. Analysis of PCR products amplified from schistosome genomic DNA using primers (indel-F, indel-R) flanking the DSBs (Fig. 2A) revealed programmed knockout (KO) efficiency, as assessed by indel (insertions-deletions)-bearing alleles resulting from the dual gRNAs, as follows: KO frequencies at GSH1 of 5.4% (range of 0.8-10.4%), 3.6% (range of 1.2-19.3%) and 12.6% (range of 4.9-19.3%) for RNP1+RNP2, RNP2+RNP3 and RNP1+RNP3, respectively (Fig. 2B). The dual RNPs mixtures induced short deletions of one to several nucleotides in length at the predicted DSB sites for sgRNA1, 2 and/or 3 (Figs. 2C-E). Mutations were not evident in amplicons from the control groups, *i.e*., wild type (no treatment) and mock treatment (not shown). The triple RNPs mixture, where the three sgRNAs shared at least six overlapping nucleotides, resulted in 23.9% KO (range of 2.4-71.9%), higher than achieved with any mixture of the dual RNPs (Fig. 2B).

The CRISPR efficiencies at each target sites varied among the RNP mixtures, as did indel profiles (Fig. S1). KO efficiency at each target resulting from dual RNPs was generally lower than for the triple RNPs. At target site of gRNA 1, estimated KO rates were 6.9% (4.1-10.4%), 13.8% (6-19.3%) and 26.5% (14-33.2%) using RNP1+RNP2, RNP1+RNP3 and RNP1+RNP2+RNP3, respectively. There were 3.9% (0.8%-5.8%), 1.8% (1.2%-2.4%) and 14.2% (2.4%-16.8%) mutations on target 2 using RNP1+RNP2, RNP2+RNP3 and RNAP1+RNP2+RNP3, respectively. Mutation efficiencies at target 3 using RNP2+RNP3 and RNP1+RNP3 were 13.9% (6%-19.3%) and 11.3% (6%-19.3%). The mutational profiles of the indels, as assessed from Sanger sequencing chromatograms of the target amplicons, were mostly deletions rather than insertions (Figs. S1B-E). Conspicuously, deletions up to 115 nt in length were identified among the biological replicates with the triple overlapping RNPs (Fig. 2E, Fig. S1E). KO efficiency was assessed using at least five independent biological replicates: Combining three overlapping sRNAs induced an aggregate mutation efficiency (23.9%) higher than that obtained with any of the dual RNPs; 5.4%, 3.6%, 12.6% (*P* ≤ 0.001, one-way ANOVA). The KO efficiency of the RNP1+RNP3 group was higher than that of either of the other dual RNPs (*P* ≤ 0.01) (Fig. 2B).

### Three overlapping guide RNAs enhanced efficiency of CRISPR knock-in

As multiple sgRNAs with overlapping sequences can enhance CRISPR/Cas9-mediated HDR efficiency ^18^ and given that triple overlapping sgRNAs performed better than dual gRNAs in initiating programmed mutation at GSH1 in eggs (Fig. 2B), we investigated KI of a reporter transgene at GSH1 with the triple overlapping sgRNA/RNPs (Figs. 3A, B). We employed the gene encoding EGFP driven by the promoter of the endogenous *S. mansoni* ubiquitin gene (Smp_335990) and its cognate terminator region as the repair template for programmed HDR (Fig. 3A). The donor template included symmetrical homology arms specific for GSH1, located on the 5’-flanking region of target site 1 and the 3’-flanking region of target site 3 (Fig. 3B). The donor template was delivered as linearized long double-strand DNA (lsDNA) of 4,451 bp in length. Aiming to optimize precise and efficient single-copy integration of the donor transgene into GSH1 by HDR, the 5’ termini of the DNA donor amplicons were chemically modified ^20^ to shield the donor template from multimerization and from integration at the DSB via the non-homologous end-joining (NHEJ) repair pathway (Fig. 3A).

**Figure 3.**
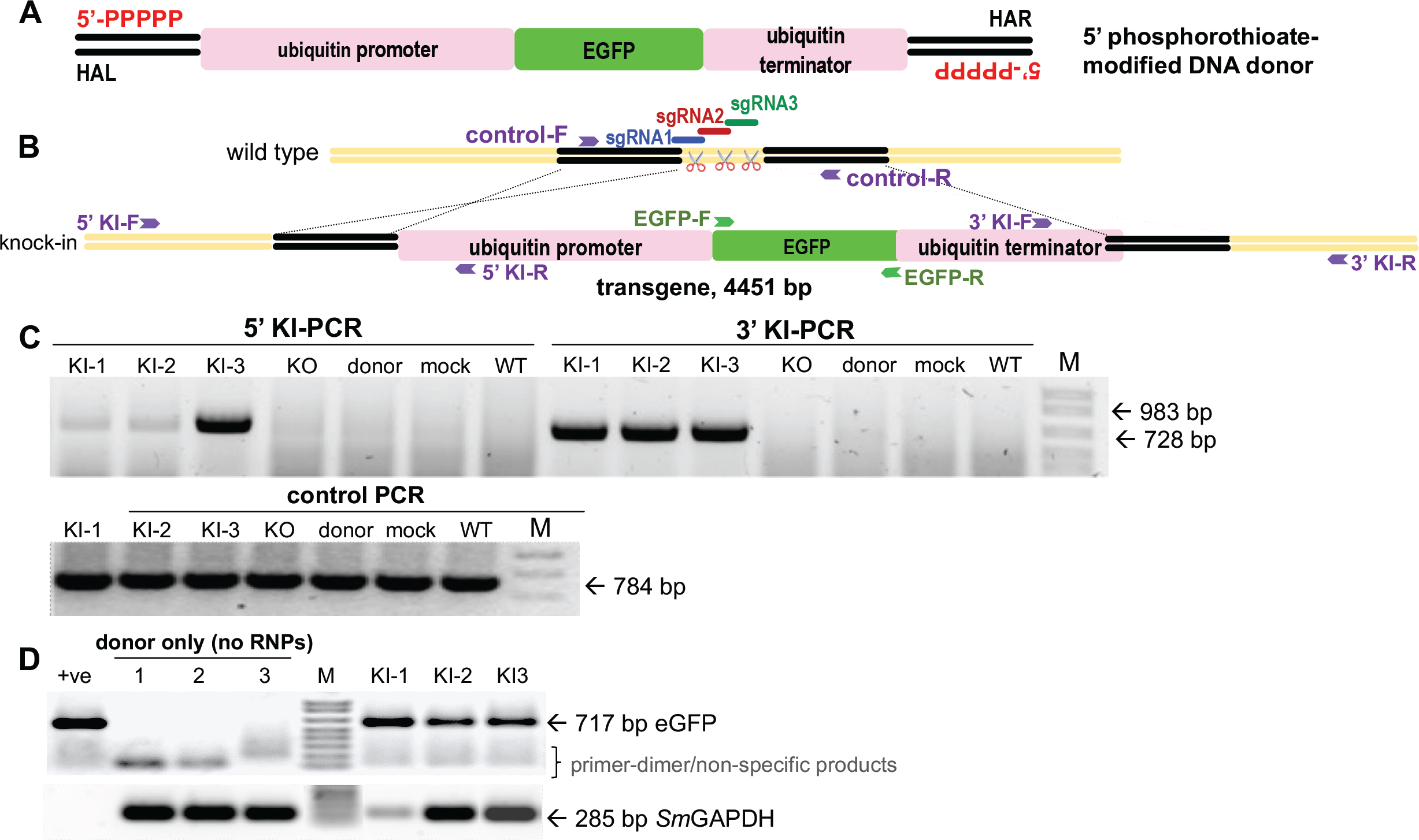
Targeted transgene insertion and expression in the schistosome egg. Programmed CRISPR/Cas9 insertion (knock-in, KI) of a donor repair template of 4.451 kb in length, encoding an EGFP transgene driven by the *S. mansoni* ubiquitin promoter and terminator. Panel A, Topology of lsDNA donor prepared from a primer pair with 5’ 5 ×-phosphorothioate modification. The donor encoded the *S. mansoni* ubiquitin promoter (pink bar) driving expression of the EGFP reporter gene (green) and ubiquitin terminator (pink) and was flanked at its termini with symmetrical 600 bp homology arms (black bars). The homology arm on the left (HAL) was situated 600 bp upstream of the position of sgRNA1 and the homology arm on the right (HAR) was sited 600 bp of downstream of the prototypic adjacent motif of sgRNA 3. Panel B, Schematic illustration of the wild type (WT) and mutant (KI) alleles after double stranded breaks programmed by the overlapping sgRNAs 1, 2 and 3. PCR primer sites are indicated by the purple arrows. Panel C. Targeted KI of the transgene detected by genomic PCR analysis using 5’KI (983 bp) or 3’KI (728 bp) specific pairs of primers. Negative controls for KI included WT, mock transfected, donor only treatment or KO (only RNPs transfection) groups, none of which had been transfected with RNPs. Lanes KI-1, KI-2 and KI-3 show amplicons from 3 independent biological replicates of programmed KI of the transgene. Other lanes show the outcome of RT-qPCRs from schistosome RNA with donor electroporation (without CRISPR materials - nuclease or guide RNAs). The integrity of the genomic DNAs was confirmed by the presence of the expected amplicon of 784 bp in all lanes of the gel labeled “control PCR” using control primer (Fig. 3B). Primer-dimer and/or non-specific PCR band(s) from donor transfected-eggs were seen ≤ 100 bp in size. Panel D. Expression of the EGFP transcript (expected amplicon size, 717 bp) as assessed by RT-qPCR following programmed KI of the transgene into GSH1. The integrity of the RNAs was assessed by analysis of transcripts of the reference gene, *Sm*GAPDH, with an expected amplicon size of 285 bp. Double-stranded DNA donor was used as the positive PCR template. Transcription of *Sm*GAPDH was seen in all treatment and control groups (lanes 1-3 and KI-1 to KI-3 in bottom panel) but not in the donor group.

At the outset, we investigated the impact of length of the homology arms (HA) by comparing the performance of the donor repair template bearing HA of increasing length of 200, 400 and 600 bp. Dual (RNP1+RNP3) and triple (RNP1+RNP2+RNP3) RNP mixtures were used in this investigation. EGFP expression was not evident in eggs electroporated with lsDNA donors with 200 and 400 bp HAs at 5 days after transfection (not shown). By contrast, we observed a few EGFP-positive eggs (∼2-3% with at least a small number of EGFP expressing cells; data from four biological replicates) with the lsDNA donor with 600 bp HA (not shown). Subsequently, we focused the investigation for EGFP expression on transfection with the donor transgene flanked by 600 bp HA using the triple RNPs (RNP1+RNP2+RNP3) and monitored EGFP expression for up to 15 days. Thereafter, on examination using spectrally resolved, confocal laser scanning microscopy (CLSM), the EGFP signals were detected in the eggs of the experimental group, which received the CRISPR materials including the lsDNA donor with 600 bp HA. EGFP signals remained until 15 days, when the experiment ended. EGFP signals were not observed in the negative control groups, although the autofluorescence characteristic of schistosome eggs was apparent ^33^. EGFP signals were also detected in the lsDNA donor control (without RNPs) for several days, indicating that extrachromosomal lsDNA expressed EGFP transiently after the transfection.

Next, we investigated the nature of the programmed KI at GSH1 by PCR-based analysis for the presence of the expected amplicons spanning the 5’ and 3’ flanks of the donor transgene, i.e., flanking the ubiquitin promoter, EGFP, and the ubiquitin terminator sequences. At the 5’-flanking region, we used a forward primer specific for the genome upstream of the 5’ HA paired with a reverse primer specific for the ubiquitin promoter (Fig. 3B). For the 3’ integration junction, a reverse primer specific for a site downstream of the 3’ terminus of the HA paired with a forward primer specific for the ubiquitin terminator was used. Fragments representing the 3’ KI and 5’ KI integration regions of 983 bp and 728 bp, respectively, were observed in the treatment groups but not in the control groups (Fig. 3C). EGFP transcripts were observed in the KI experimental group, although some variability in transcript abundance among the biological replicates was seen based on the signals obtained for *Sm*GAPDH, which served as the reference gene (Fig. 3D).

### Reporter transgene expression in edited schistosome eggs

EGFP positivity and intensity were quantified using spectral laser scanning CLSM ^33^. Active transgene expression was confirmed within miracidia developing inside transfected eggs (Figs. 4A, B). EGFP appeared to be expressed by numerous diverse cells throughout the developing larvae, whereas morphological malformation was not observed in transgenic eggs and their enclosed larvae. More intense EGFP fluorescence was consistently recorded and quantified at 509 nm in eggs from the experimental treatment group (Figs. 4B1, 2) than the mock control eggs, and in eggs transfected solely with donor template (Figs. 4A1, 2). Subsequently, on day 15 following transfection with the repair template in the presence or absence of the RNPs mixture, we quantified EGFP intensity in eggs that contained a miracidium by normalization with the EGFP signal from lsDNA only transfected eggs. Fluorescence intensity differed markedly between these two groups: by 15 days after transfection, 75% of miracidia in eggs from the experimental group (RNPs/lsDNA) emitted EGFP fluorescence, whereas 25% of eggs containing a miracidium transfected with the donor lsDNA repair template only emitted EGFP (Figs. 4C, D) (*P* ≤ 0.001; *n* = 402 eggs in the experimental group, *n* = 397 eggs in the lsDNA only group (Fig. 4E), collected from four independent biological replicates).

**Figure 4.**
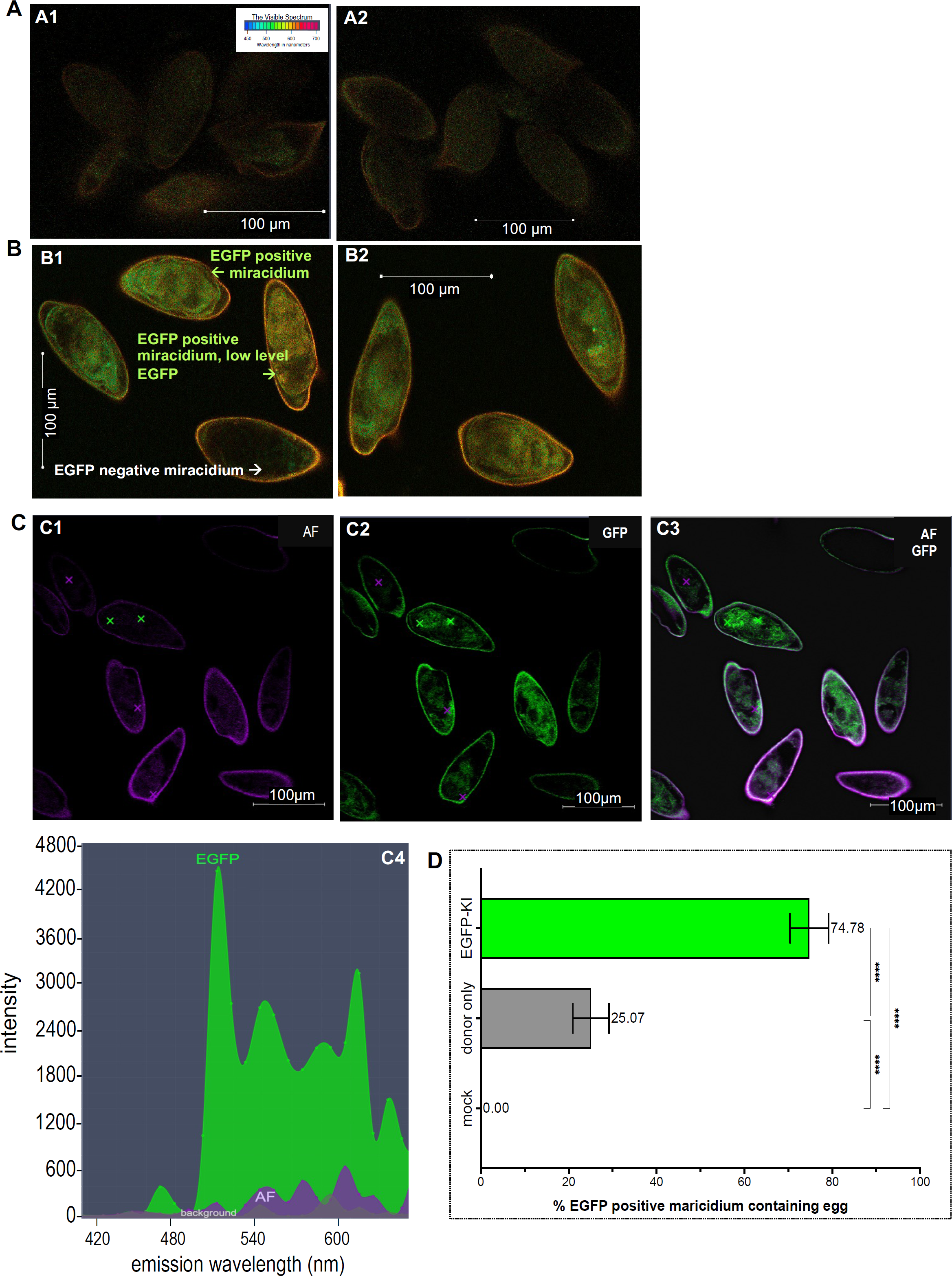
Higher numbers of EGFP-positive eggs following programmed knock-in at GSH1 as assessed by spectral image analysis. Confocal laser scanning micrographs: Panel A. Eggs exhibiting background signal (autofluorescence; AF) from the control group, *i.e.* eggs transfected only with the lsDNA donor repair template; A1, A2, representative images from biological replicates. Panel B. Eggs expressing EGFP in the experimental group transfected with RNPs and the donor repair template; B1, B2, representative images from two biological replicates. Many eggs expressed EGFP with the broad range in intensity of fluorescence ranging from higher intensity (green arrow) and lower levels (yellow arrow) following programmed KI. Eggs expressed EGFP observed from day 5 until day 15 when the experiment was terminated and taken the micrographs. Panel C, Linear unmixing analysis revealed EGFP fluorescence in miracidia within the eggshell, as shown in these representative micrographs, C1, C2 and C3; C4, plot showing the emission spectrum of EGFP (green curve) with a peak signal at 509 nm (× in bright green indicates the EGFP signal, while the × in purple color indicates the AF inside the egg), and, also showing the autofluorescence background (purple curve). Panel D, To assess expression of EGFP, eggs were counted (scored) as EGFP-positive when >20% of the surface area of the miracidium (within the eggshell) was EGFP positive; representative EGFP-positive eggs are shown in panels C2 and C3 (green eggs). On day 15 following transfection, eggs were scored and counted for EGFP positivity in each of 4 independent biological replicates (∼100 eggs from each replicate) of KI (RNPs + lsDNA) and lsDNA only eggs. About 25% of the eggs in the lsDNA only control group were scored as EGFP-positive (range, 19 to 32%), whereas 75% of eggs the KI group expressed EGFP (range, 68 to 79%); *P* <0.001; two-tailed *t* = 69.87, *df* = 142; difference between means (EGFP-KI-only donor) ± SEM, 49.7 ± 0.7, 95% CI, 48.3 to 51.1. There was marked decrease of background autofluorescence in eggs from WT and mock-transfection groups after the linear unmixing normalization of the fluorescence spectrum.

In addition, we scored the intensity of fluorescence at 509 nm ^33^, the emission wavelength for EGFP, as shown in Fig. 4, panels C2, C3 and D (green curve). To this end, we subtracted the signal at 509 nm from the autofluorescence spectrum, which originated from the eggshell. The EGFP-specific signal in the control lsDNA donor repair template treatment group, mean = 1,290 arbitrary units (AU) ^34^ (range, 856-1,1713) was significantly lower than the experimental group transfected with the triple, overlapping guide RNPs mixture in the presence of the donor repair template, 6,905 AU (range, 4,973 to 8,963) (*P* ≤0.001) (supplement Fig. 2). Moreover, emission of EGFP was not detected in the control groups, *i.e.* mock-transfected and WT eggs (not shown). Diverse cells and tissues of the developing miracidium expressed the fluorescence reporter gene, thus EGFP expression appeared not be restricted to specific cells (Figs. 4B, C).

### Impact on egg viability of delivery of CRISPR materials by electroporation

During the investigation, we also examined delivery of CRISPR materials using electroporation of the schistosome egg, an approach originally described for transfection the schistosomulum stage of *S. mansoni*^35^. At the outset, electroporation voltage was investigated, using a single pulse of 20 ms duration of 125, 150, 200 and 250 volts (V) to deliver the RNPs and donor repair template into the eggs. Thereafter, the eggs were cultured and miracidial hatching assessed seven days later. Survival and/or larval growth inside the egg in the 125V treatment group was not significantly affected; the rates of miracidial hatching were 26.6±3.2% and 31.9±2.6% from non-electroporated group (Fig. S3A; supplemental information). By contrast, increasing the voltage negatively impacted hatching of miracidia from the eggs: 150V, 22.8±2.2%; 200V, 11.4±1.2%; 250 V, 3.9±2.0%, respectively (*P* < 0.001, two-way ANOVA) (Fig. S3A). Last, we investigated the impact of the CRISPR materials in addition to voltage. Using electroporation at 125V, we monitored hatching in two biological replicates. In the first, 41.2±2.1, 39.5±1.5 and 40.9±1.9% (mean ±SE) of miracidia hatched from the wild type, donor transfection only and experimental (EGFP knock-in) groups, and in the second, 59.1±2.4%, 60.5%±0.6 and 60.1±1% from wild type, donor transfection only and EGFP knock-in groups, respectively (Figs. S3B, C).

## Discussion

To advance functional genomics for helminths, we identified four potential GSH sites in *S. mansoni*, optimized conditions for delivery and structure of transgene cargo, inserted the reporter transgene into the most qualified intergenic genome safe harbor, GSH1, by programmed CRISPR/Cas9 homology directed repair. We confirmed integration of the transgene by amplicon sequencing as well as EGFP reporter activity using RT-PCR and CLSM analyses. Our approach for programmed editing in this helminth involved electroporation based delivery to the schistosome egg of RNPs with overlapping guide RNAs, in the presence of phosphorothioate-modified, double-stranded donor targeting at GSH1. The procedure yielded 24% editing efficiency that was accompanied by transgene activity in 75% of miracidia in the genome-edited schistosome eggs. The donor dsDNA encoded a reporter construct consisting of the EGFP gene driven by the schistosome ubiquitin promotor and terminator. Furthermore, clear EGFP signals indicated the suitability of the regulatory elements of the ubiquitin gene to induce transgene expression.

This methodical approach provides a tractable path towards transgenic helminths using homology-directed repair-catalyzed transgene insertion. Our criteria to predict GSH included location in euchromatin to avoid silencing of the transgene, unique genome-target sequence to minimize off-target events, avoidance of lncRNA encoding genes, presence of epigenetic marks for open chromatin structure, and the absence of epigenetic marks indicating heterochromatin. We termed the intergenic sites GSH1, -2, -3 and -4, which were located on chromosomes 2 and 3. (*S. mansoni* has seven pairs of autosomes and one pair of sex chromosomes, Z and W, with the female schistosome being the heterogametic sex^36^). In addition, we assessed the GSH1 locus for CRISPR/Cas9 integration, gene editing and over expression of EGFP. We edited GSH1 using ribonucleoprotein complexes of Cas9 endonuclease with multiple, overlapping guide RNAs. Overlapping CRISPR/Cas9 target sites were selected using the CHOPCHOP web-based tool. RNPs of triple overlapping guides delivered significantly higher CRISPR/Cas9 efficiency than RNPs of dual guides, and longer length deletion mutations. In addition, efficient HDR was obtained using a combination of multiple target site overlapping RNPs programmed to cleave GSH1 in the presence of a repair template protected by chemical modifications. Our approach successfully inserted a long (4,551 bp) doubled stranded DNA donor template at GSH1. This outcome aligns with reports in cell lines and rodents involving overlapping gRNAs, where deletions close to the targeted mutation enhanced the efficiency of HDR ^18, 37^. Overlapping guide RNAs rather than simply deploying multiple guides may be more efficient for gene knockout in *S. mansoni* given recent findings involving CRISPR interference that compared both single and multiple guide RNAs ^18, 19, 38^.

Accordingly, GSH1 represents a promising safe harbor region for *S. mansoni*. Notably, 75% of mature eggs exhibited EGFP fluorescence in the miracidium developing within the eggshell, and significantly more fluorescence than seen in the control eggs transfected with donor template but not with the RNPs (∼5%). EGFP signals were not present in the control, untreated wild type eggs, which by ∼10 days following transfection exhibited minimal background fluorescence. Our approach to evade the autofluorescence emitted by schistosome eggs, which can confound detection of EGFP, used CLSM spectral imaging and linear unmixing ^39, 40^, a new approach to facilitate quantification of EGFP-specific emission to resolve overlap between the EGFP and endogenous fluorophores in schistosomes. Freshly prepared eggs isolated from livers of *S. mansoni-*infected mice (∼ 42 days after infection) were transfected using electroporation of two or three RNPs and the donor transgene. Such a preparation of schistosome eggs includes eggs displaying a spectrum of development – from newly laid eggs containing an immature miracidium, maturing eggs, eggs containing the fully developed miracidium (miracidia movement inside the egg is apparent), and likely some dead eggs as well^41^. Notably, however, the entry of active CRISPR materials and donor transgene into cells of each egg and each developmental stage cannot be predicted. The suitability of this approach for introducing Cas9 RNPs into LEs has been demonstrated by RNP tracking analysis^42^. Indeed, the outcome would be stochastic: not every egg would be expected to receive the full complement of RNPs and donor. Accordingly, eggs exhibiting minimal EGFP may have not been transfected as efficiently as eggs with stronger EGFP fluorescence.

Fluorescence throughout miracidial tissues was achieved using EGFP driven by the schistosomal ubiquitin promoter and terminator, emphasizing its ubiquitous activity as predicted by transcriptome analyses^43–45^. This outcome confirmed reporter-gene activity under the control of these ubiquitin elements and demonstrated the accessibility of GSH1 for the transcriptional machinery after programmed gene editing. The findings also revealed the feasibility of selection at the microscopic level, which would enable hand-picking of reporter gene-positive miracidia for snail infection to complete the life cycle. Following snail infection, fluorescing cercariae - or in case of mono-miracidial infections also reporter gene PCR-positive cercariae - could be selected for infection of laboratory rodents to generate heritably transgenic worms. In future approaches, GSH1 may alternatively be used as a locus to integrate other transgenes, e.g., antibiotic resistance gene(s), which would enable drug selection at the stage of embryogenesis or during the intermediate host stage in the snail. Here, oxamniquine is a suitable candidate drug^42^. In case mono-miracidial infections with reporter gene-positive miracidia would be performed, additional selection manipulations (microscopic or PCR-based) could be performed at the stage of the clonal cercariae obtained by this approach. Thereafter, reporter gene-positive female and male cercariae, in which gender can be confirmed by quantitative PCR^46^, could be mixed for infection of the mammalian host as a basis for genetic crosses^47^.

We focused on GSH1 because there were apparent limitations to progress with the other three prospective GSHs, GSH2-4. The CHOPCHOP software predicted only a single CRISPR/Cas9 target in each of these three GSHs and, moreover, the site predicted in GSH4 was not specific and showed potential off-target hits elsewhere in the genome. For GSH3, CHOPCHOP predicted only low, <35%, editing efficiency. Overall, locating a Cas9 PAM, i.e., the NGG motif ^48^, is constrained by the AT-rich nature of the genome of this blood fluke^49^. Moreover, since this investigation deployed multiple, overlapping gRNAs^18^ to facilitate homology directed insertion of transgene of ∼4.5 kb in length, we ranked GSH1 as the most qualified for our purposes because multiple PAMs were present, along with the absence of off-targets and donor homology arms did not exhibit DNA complexity and hairpins. Nonetheless, comparisons of the utility of all four GSH including with additional nucleases such as Cas12a ^17^ and base editors ^50^ could be considered in the future. Although a distance > 2 kb from known genes was one of our criteria for GSH in *S. mansoni*, intragenic sites rather than intergenic GSH nonetheless may have expedient attributes for functional genomics where partial loss of fitness may be less consequential, e.g., for *in vitro*-focused studies with *S. mansoni*. In experimental human gene therapy ^24^, the most widely targeted sites are intragenic safe harbors, *CCR5*, *AAVSI*, and *Rosa26*, and despite their gene-rich nature, they have been targeted with therapeutic gene cargo, including the insertion of *FANCA* at *AAVS1* in CD34^+^ hematopoietic progenitors from Fanconi anemia cases ^51^. Yet, intergenic sites are inherently safer given coding regions or other elements are not disrupted. Recently, additional sites, named *Rogi 1* and *Rogi 2*, located in intergenic regions, conform with additional criteria focusing on enhancement of safety for use in human gene therapy, including location distant from endogenous genes, lncRNAs, enhancers, and telomeres ^28^.

These findings are consequential in that they advance functional genomics for a hitherto unmet challenge to manipulate a pathogen of global public health significance. They confirm that transgenes can be inserted into a predicted GSH to endow individual stages or populations of these pathogens with novel functions, with broad potential for basic and translational studies ^52–54^. Whereas this report deals with somatic transgenesis of the schistosome larva, use of the same approach for transfection of the newly laid egg (NLE) of *S. mansoni*, a stage that at its origin includes a single zygote (surrounded by vitelline yolk cells). The NLE might represent a window to the germline, and the hypothesized accessibility of its zygote may facilitate complete transformation to derive lines of transgenic parasites carrying gain- or loss-of-function mutations. In addition, the gene editing methods developed here can be adapted for knockout approaches of other genes of interest, in schistosomes, and likely other platyhelminths, for which genome sequences are available to be analyzed for GSHs. The information presented provided novel insights into efficient transgenesis and forward genetics for *S. mansoni* for this and prospectively further parasitic (and free-living) helminths.

## Acknowledgments

Schistosome-infected mice were provided by the Schistosomiasis Resource Center of Biomedical Research Institute, Rockville, MD through NIH-NIAID contract HHSN272201700014I for distribution through BEI Resources. This research was funded in part by the Wellcome Trust grant 107475/Z/15/Z (Flatworm Functional Genomics Initiative, PI, Karl F. Hoffmann). For the purpose of open access, the author has applied a CC BY public copyright licence to any Author Accepted Manuscript version arising from this submission. This study is set within the framework of the « Laboratoire d’Excellence (LabEx) » TULIP (ANR-10-LABX-41) with the support of LabEx CeMEB, an ANR « Investments d’avenir » program (ANR-10-LABX-04-01) and the Environmental Epigenomics Core Service at IHPE (Interactions Hôtes Pathogènes Environnement). PJB gratefully thanks LabEx CeMEB for support as a short term visiting scientist at IHPE, Université Perpignan via Domitia. Award number RR025565 (PI, Anastas Popratiloff) from the NIH Office of Research Infrastructure’s S10 program supported the purchase of the Zeiss 710 confocal microscope.

## Author contributions

Conceptualization by WI, CG, CGG and PJB. CC and CG software predicted and identified gene safe harbors. MFM and TQ cloned and analyzed the active promoter/terminator. WI proceeded the methodology, validation and mutation analysis of gene editing for both knock-out and knock-in. RR, PW, and WB optimized and investigated the donors used for knock-in condition. VM, AM, LM, SS, and MMK provided the parasite resources such as schistosome life cycle maintenance and parasite egg preparation. MFM, AP, and WI visualized the reporter gene and analyzed confocal microscopic visualization and analysis: MFM, AP, WI. WI, CG, CGG and PJB are supervision of the study. WI, CG, CGG, and PJB drafted, reviewed, and edited the manuscript.

## Data and materials availability

All data are available in the main text or the supplementary materials. The nucleotide sequence reads are available at the NIH Sequence Read Archive, BioProject PRJNA919068, accession numbers SRX18957908-18957932. BED files from the bioinformatics analysis are available on Zenodo, https://zenodo.org/record/7602535#.ZAIgqBPMLIM.

## STAR Methods

### Computational search for gene safe harbors in *Schistosoma mansoni*

We undertook a genome analysis focusing of intergenic (gene-free) regions to identify prospective GSHs, using similar approaches as those used on the human genome ^28^. We aimed to locate a GSH, a site that would facilitate stable expression of the integrated transgene free of interference from the host genome and which, in parallel, integrates and transcribes the transgene without negative consequences or loss of fitness for the host cell. The search for GSHs deployed included several criteria, First, its location should be adjacent to peaks of H3K4me3, a histone modification associated with euchromatin and transcription start sites ^55^. Second, it should not be near or not containing H3K27me3 in any developmental stage, a histone mark associated with heterochromatin ^55^. Third, as the schistosome genome contains highly repetitive elements ^49^, the GSH site should be located in a unique tract of the genome sequence. Fourth, it should reside in open, euchromatic chromatin accessible to Tn5 transposase as assessed from ATAC-sequencing, which provides a positive display of transposase integration events ^56^; consequently, safe harbor candidate regions should deliver an ATAC-sequence signal. Fifth, in the vicinity of known HIV integration sites, given that HIV integrates preferentially into euchromatin in human cells ^57^, we anticipated that HIV integration into the schistosome genome may likewise indicate a region of euchromatin (Fig. 1A) ^58^.

To predict loci conforming to the criteria, pooled ChIP-seq data for H3K4me3 and K3K27me2 from previous studies were aligned against on *S. mansoni* genome data (version 9 on the date of analysis). ATAC-seq was performed as described with modification ^59^. Peakcalls of ChIP-seq and ATAC-Seq were done with ChromstaR ^21, 28, 55, 56^ and stored as Bed files. Bed files were used to identify the presence of H3K4me3 and absence of H3K27me3 in adults, miracidia, *in vitro* sporocysts, cercariae and *in vitro* schistosomula with bedtools intersect. Thereafter, ATAC-seq data from adult male and adult female worms (two replicates each) were intersected to find common ATAC-positive regions. H3K4me3-only (H3K27me3-absent) common to all stages and ATAC signals were intersected to find common regions. Next, the HIV integration sites were identified by using data from ERR33833.8. Reads were mapped to the lentivirus genome (HIV-1 vector pNL-3, accession AF324493.2) using Bowtie2 with default parameters. Paired reads were extracted where one end mapped to HIV and the other end mapped to schistosome genome at a unique location. Genes from the BED files above that located ≤ 11 kb HIV-1 integration sites were identified with bedtools closestbed. Gene expression data for these genes were obtained using the metanalysis tool, https://meta.schisto.xyz/analysis/, of Lu and Berriman ^44^.

Computational searches that addressed these criteria predicted, *a priori*, gene free (intergenic)-GSH (Fig. 1), given that transgene integration into an existing gene could diminish fitness of the genetically modified cell ^23, 24^. We defined genes as protein coding sequences and sequences coding for long non-coding RNA (lncRNA). In view of our goal to use CRISPR/Cas mediated-HDR to insert the transgene, we searched preferentially for unique sequences, to obviate off-target gene modification, and excluded gene free-regions composed of repetitive sequences. Those unique sequences were also annotated outside lncRNA, regions beyond putative promotors that we deemed as 2 kb upstream of the transcription termination site (TTS), and the regions close to peaks of H3K4me2 in all parasite stages which never contained H3K27me3. The regions overlapping with ATAC-seq positive sites with ≤11 kb distance from HIV integration sites also were included. (The HIV-1 genome is ∼10 kb in length.) BEDtools were used to delimit 2 kb upstream regions (FlankBed). Annotations of 16,583 lncRNA were pooled from http://verjolab.usp.br/public/schMan/schMan3/macielEtAl2019/files/macielEtAt2019.bed12 ^60^. Repeats were masked with RepeatMasker V4.1.0 using a specific repeat library produced with RepeatModeler2 V2.0.1 and stored as a GFF file. BED files with coordinates outside these annotations were generated by BedTools complementBed. BedTools Multiple Intersect was used to identify regions that are common to unique regions (complement of repeatmasker), intergenic regions, ≥ 2 kb upstream and outside of lncRNA. Regions which a length ≥ 100 bp were retained. These regions were intersected with merged H3K4me3-only common to all developmental stages and ATAC signals (euchromatic signal). BedTools ClosestBed was used to determine distance to the nearest integrated HIV provirus.

### Schistosome eggs

Mice (female, Swiss Webster) infected with *S. mansoni* were obtained from the Schistosomiasis Resource Center (Biomedical Research Institute, Rockville, MD) within seven days of infection by cercariae (180 cercariae/mouse/percutaneous infection). The mice were housed at the Animal Research Facility of George Washington University, which is accredited by the American Association for Accreditation of Laboratory Animal Care (AAALAC no. 000347) and has the Animal Welfare Assurance on file with the National Institutes of Health, Office of Laboratory Animal Welfare, OLAW assurance number A3205. All procedures employed were consistent with the Guide for the Care and Use of Laboratory Animals. The Institutional Animal Care and Use Committee of the George Washington University approved the protocol used for maintenance of mice and recovery of schistosomes.

Mice were euthanized six to seven wk after infection, after which schistosomes were recovered by portal vein perfusion with 150 mM NaCl, 15mM sodium citrate, pH 7.0. The liver was resected, homogenized with a. tissue blender, and the homogenate incubated with collagenase at 37°C for 18 h. Thereafter, schistosome eggs from the digested livers were recovered by Percoll gradient centrifugation, as described ^61^. Eggs isolated from livers, termed “liver eggs”, LE ^62^, were cultured in 20% inactivated bovine serum, 2% antibiotic/antimycotic at 5% CO_2_, 37°C overnight before being subjected to transfection with the CRISPR materials.

### Guide RNAs, ribonucleoprotein complexes

Here, we focused on GSH1, located on *S. mansoni* chromosome 3; 13231955-13233370 (Fig. 1D), an intergenic safe harbor site of 1,416 nt, the longest in length of the four GSH (Figs. 1B-D). Guide RNAs (gRNA) for GSH1 were designed with the assistance of the CHOPCHOP ^30, 31, 63^ tools, using the annotated *S. mansoni* genome ^64^, to predict target sites, off-targets, and efficiency of CRISPR/Cas9 programmed cleavage. CHOPCHOP predicted three overlapping guide RNAs targeting three cleavage sites withinGSH1 with the three DSBs at six to 12 nt apart from each other. All three overlapping gRNAs lacked off-target sites and lacked self-complementarity. The three gRNAs were located on the forward strand of GSH1 at nucleotide positions 605-624, 617-636, and 623-642, respectively (Fig. 2A). We termed the predicted DSBs as target 1, target 2 and target 3. Synthetic guide RNAs (sgRNA), specifically Alt-R CRISPR-Cas9 sgRNA chemically modified to enhance functional stability, and recombinant *Streptococcus pyogenes* Cas9 nuclease, specifically Alt-R HiFi Cas9, which includes nuclear localization sequences (NLS), were sourced from Integrated DNA Technologies, Inc. (IDT, Coralville, IA). Each ribonucleoprotein complex (RNP) was complexed in the separate tube, with Cas9 and a single sgRNA at 1:1 ratio in 25 µl Opti-MEM; 10 μl of 1 μg/μl sgRNA (Opti-MEM as diluent) was mixed with 10 μl of 1 μg/μl Cas9 (in Opti-MEM) by gentle pipetting and incubated for 10 min at 23°C to allow the RNP to assemble.

### Doubled stranded DNA donor

A plasmid vector, pUC-Ubi-EGFP-ubi, was constructed by chemical synthesis of the donor transgene and its ligation into pUC (Azenta Life Sciences, Chelmsford, MA). The inserted sequence included homology arms of 600 bp length corresponding to GSH1 at 22-621 nt (5’-homology arm) and 640-1239 nt (3’-homology arm), respectively, flanking the in-frame expression cassette of the *S. mansoni* ubiquitin promoter (2,056 bp), EGFP (717 bp), and the ubiquitin terminator (578 bp). Plasmid DNA was amplified using Phusion High-Fidelity DNA Polymerase (New England Bio-Labs, Ipswich, MA, cat no. M0530) with primers specific for the 5’ and 3’ termini of the homology arms of 200, 400 or 600 bp in length (Table S1). Thus, three forms of the donor transgene DNA were prepared, with homology arms of increasing length – 200, 400 and 600 bp. The primers employed to amplify the transgene DNA from the pUC-Ubi-EGFP-ubi plasmid were 5×phosphorothioate-modified to enhance stability of the amplified DNA. (5’-modified long dsDNA donor (lsDNA) enhances HDR and favors efficient single-copy integration by its retention of a monomeric conformation ^20^. (Fig. 3A). PCRs were carried out in reaction volumes of 50 μl in 200 μM dNTPs, 0.5μM of each primer, 100 ng pUC-Ubi-EGFP-ubi, 3% DMSO and 1 unit of Phusion DNA polymerase, with thermocycling of 98°C, 30 sec, 30 cycles of 98°C, 10 sec, 55°C, 30 sec, 72°C, 3 min, and final extension at 72°C, 10 min. Amplificons were isolated using the NucleoSpin Gel and PCR Cleanup and gel extraction kit (Takara, San Jose, CA, cat no. 740609), eluted in 30 μl nuclease-free water, and the long stranded (ls) DNA donor transgene stored at -20°C until used.

### Transfection of schistosome eggs

Ten thousand eggs (LE) of *S. mansoni* were washed three times with chilled (4°C) 1×PBS before transfer into a chilled electroporation cuvette (4 mm electrode gap, BTX, Holliston, MA) with Opti-MEM as the electroporation medium. Each 25 μl of RNP along with the lsDNA donor were immediately dispensed into the cuvette containing the schistosome eggs, to a total cuvette volume of 300 μl with Opti-MEM: specifically, for the dual guide RNA/RNPs, group 1) 25 μl RNP1+25 μl RNP2, group 2) 25 μl RNP2+25 μl RNP3, and group 3) 25 μl RNP1+25 μl RNP3. In groups with the lsDNA, 10 μg of this donor DNA was added into the cuvette before bringing the final volume to 300 μl/cuvette. Transfection of schistosome eggs with CRISPR materials was undertaken using square wave electroporation (Electro SquarePorator ECM 830, BTX). Using a single pulse of 125 volts for 20 ms^16, 17, 65^ was confirmed as optimal for use in this study based on analysis using higher voltages; viability as indicated by % miracidial hatching decreased progressively as voltage increased to 150, 200 and 250 V (Fig.S3). The transfected eggs were transferred to culture medium, as above.

### Nucleic acids

To recover genomic DNA and total RNA, eggs from each replicate were triturated in ∼100 μl DNA/RNA Shield solution (Zymo Research, cat no. R1100, Irvine, CA) using a motor-driven homogenizer fitted with a disposable pestle and collection tube (Biometer II, Bunkyo-ku, Tokyo, Japan). DNA was isolated from 50% of the homogenate, and RNA from the remainder. 250 μl DNAzol^®^ ES (Molecular Research Center, Inc., Cincinnati, OH) was dispensed into the homogenate, and DNA recovered according to the manufacturer’s protocol. Total RNA was extracted from the homogenate by adding 250 μl RNAzol RT (Molecular Research Center, Inc.). Yields and purity were assessed quantified by spectrophotometry (NanoDrop One Spectrophotometer, Thermo Fisher Scientific), based on the ratios of absorbance at 260/280 and at 260/230 nm ^66^.

### Analysis of CRISPR on-target efficiency

Amplicons of GSH1 spanning the programed DSBs were obtained using population genomic DNA (above) and primers termed ‘indel-F and indel-R primers’ that cover the region flanking expected double strand break of all the CRISPR target sites (Fig. 2A, Table S1). Amplification products were purified (NucleoSpin Gel and PCR Cleanup and gel extraction kit, Takara; cat no. 740609) and the nucleotide sequences determined by Sanger cycle sequencing (Azenta Life Sciences, South Plainfield, NJ). Chromatograms of the sequence traces of experimental and control group(s) was compared using DECODR ^32^ at default parameters with single and multiple CRISPR target analysis (Figs. 2B-F, S1A). The pattern of indels in was confirmed by TIDE analysis of Sanger sequence reads ^67^ (Figs. S1B-E).

### Detection of integration of the transgene into the schistosome genome

Integration of the donor transgene at GSH1 was analyzed by PCR, using GoTaq^®^ G2 DNA polymerase (Promega, Madison, WI) and two pairs of primers; one primer located on the GSH1 using specific primers upstream (5′ KI-F) or downstream (3′ KI-R) of the homology arms paired with primers specific for the transgene (5′ KI-R or 3’ KI-F) (Fig. 2B, Table S1), as described ^17^. The PCR cycling conditions were 95°C, 2 min, 40 cycles 94°C, 15 sec, 58°C 30 sec, 72°C, 60 sec, and the amplification products were separated by electrophoresis and stained with ethidium bromide. The expected product sizes for the 5 ′ and 3′ integration site-specific amplicons were 728 and 983 bp, respectively, and the amplification control (amplified by indel-F and indel-R primers), expected product size, 764 bp (Fig. 2C).

### Quantification of transgene expression in schistosome eggs

To examine the mRNA expression of EGFP, total RNAs extracted from LE were exposed to DNase to eliminate residual genomic DNAs and donor lsDNA donor were transcribed into cDNA using the Maxima First Strand cDNA synthesis kit with DNase (Thermo Fisher Scientific). The qPCR was performed using the GoTaq^®^ G2 DNA polymerase (cat no. M7841, Promega, Madison, WI) with the specific primers; EGFP-F and EGFP-R (Fig. 3b, Table S1) with expected amplicon at 717 bp. *S. mansoni* GAPDH (Smp_056970) served as the reference gene. The specific primer for GAPDH Table S1) expected amplicon of 285 bp in length. PCR cycling conditions: 95°C, 2 min, 25 cycles 94°C, 15 sec, 58°C, 30 sec, 72°C, 30 sec, after which amplification products were examined, as above.

### Quantification of fluorescence by spectral imaging and linear unmixing

Spectral and spatial distribution of EGFP fluorescence were assessed using confocal laser scanning microscopy, using a Zeiss LSM710 Meta detector fitted Axiovert 200 (Carl Zeiss, Jena, Germany). Images were collected with the W Plan-Apochromat 20×/1.0 NA water immersion objective. Spectroscopic measurements were performed in response to excitation by 458 nm (16.5 μW) Argon laser line and 633 nm He/Ne laser line (Lasos Lasertechnik, Jena, Germany), which were used for focus and transmission mode imaging. Emission was detected with a spectral META detector at 16 channels, scanning at 477 nm through 638 nm, simultaneously. A hurdle for the detection of EGFP using fluorescence microscopy is presented by the autofluorescence known to originate from the schistosome eggshell ^68–70^, with vitelline cells the likely origin of this emission ^71^. To surmount this hurdle, the EGFP spectrum emitted from within each eggshell was investigated by selecting the area to be examined, specifically the entire miracidium, and collecting multispectral images of the miracidium within the eggshell using the LSM Image Examiner. The images collected were assessed for EGFP by subtracting regions emitting autofluorescence from the fluorescence signal collected for entire surface area of the egg: Specifically, the total EGFP intensity at 509 nm ^71^ was calculated using the Zeiss Zen (black edition) software module from ∼400 eggs (∼100 eggs from each of four biological replicates) in each of the control and experimental groups.

## Supporting information

**Figure S1.**
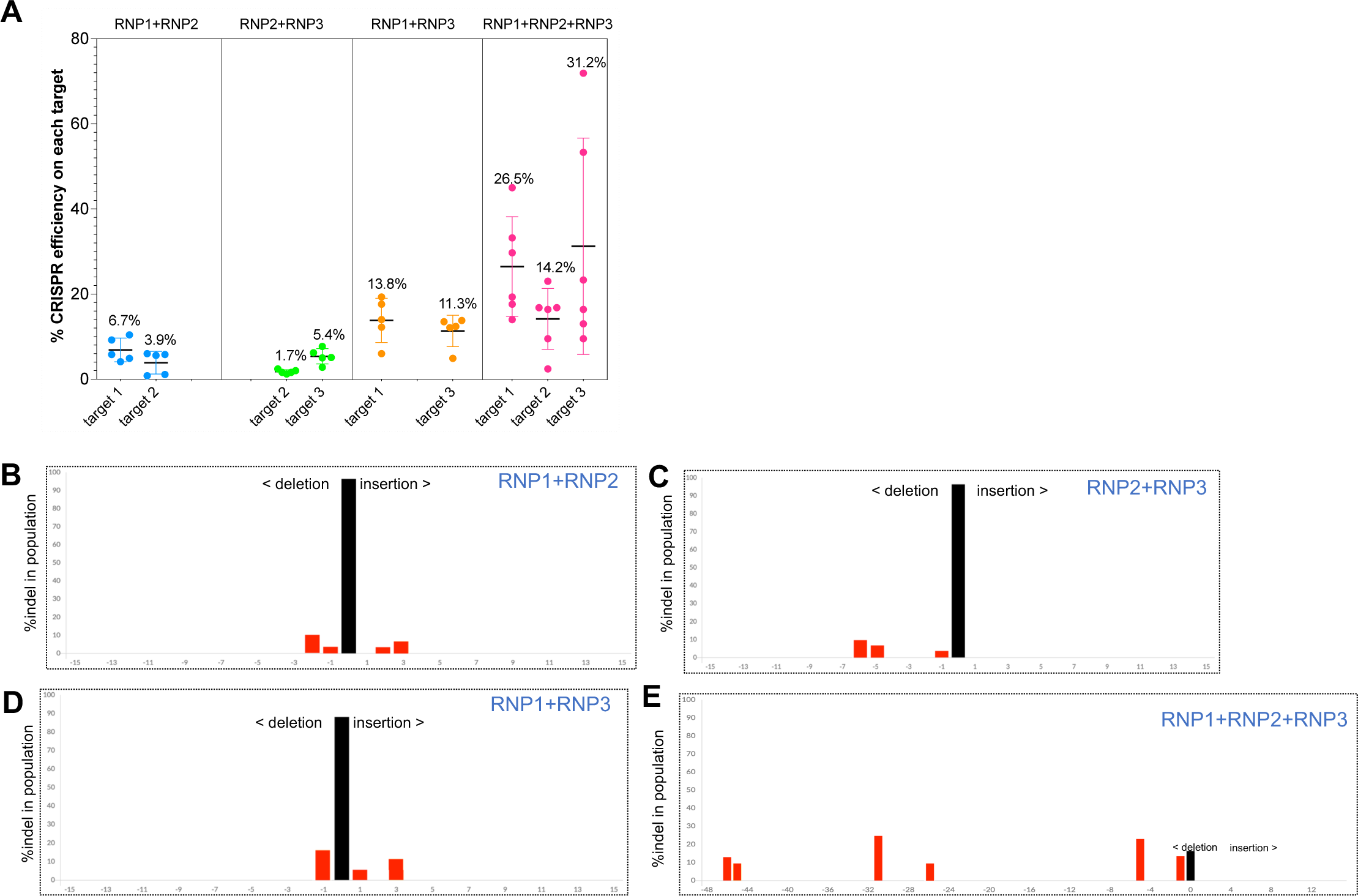
Panel A, CRISPR efficiency (percentage) on individual target sites as estimated using the Deconvolution of Complex DNA Repair (DECODR) algorithm using distance from dual gRNAs; RNP1+RNP2 (blue dots), RNP2+RNP3 (green), RNP1+RNP3 (yellow) and triple gRNAs; RNP1+RNP2+RNP3 (pink dots), plotted from 5-6 independent biological replicates. The % indels on each target varied among RNP combinations:| transfection with triple RNPs lead to highest CRISPR efficiencies on each target site 1, 2 and 3, 26.5%, 14.2% and 31.2%, respectively, compared with the dual RNP groups. The % indels resulting from dual RNPs on target 1 were 6.7% and 13.8%; target 2, 3.9% and 1.7%, and target 3, 5.4% and 11.3%. Panels B-E, examples of indel patterns resulting from the RNPs. DECODR software plots confirmed the indel profiles and were similar to those obtained with TIDE (Fig. S1B - E).

**Figure S2.**
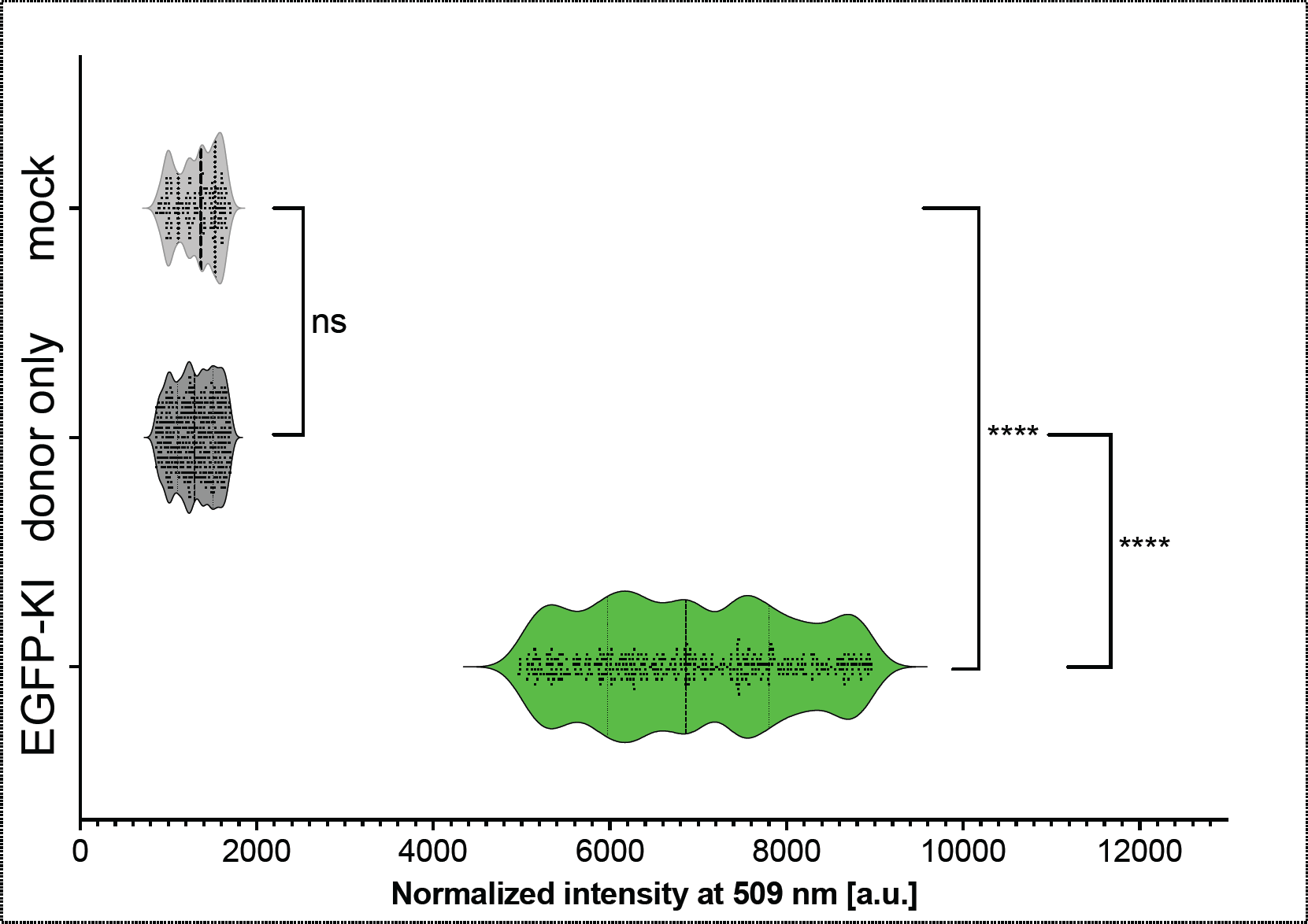
Emission spectral intensity for eggs, scanned from 477 to638 nm, normalized fluorescence spectral intensity from control eggs (transfected with donor repair template) exhibiting higher intensity than autofluorescence; these eggs were also scored as EGFP-positive, and with a normalized EGFP intensity mean, 1290 au (range, 856-1713); experimental group, normalized-EGFP intensity, mean 6905 au (range 4972 – 8963); *P* < 0.001, unpaired *t*-test, *n* = 402; difference between means of experimental and control group eggs ± SEM, 5651 ± 57.4, 95% CI, 5502 to 5728).

**Figure S3.**
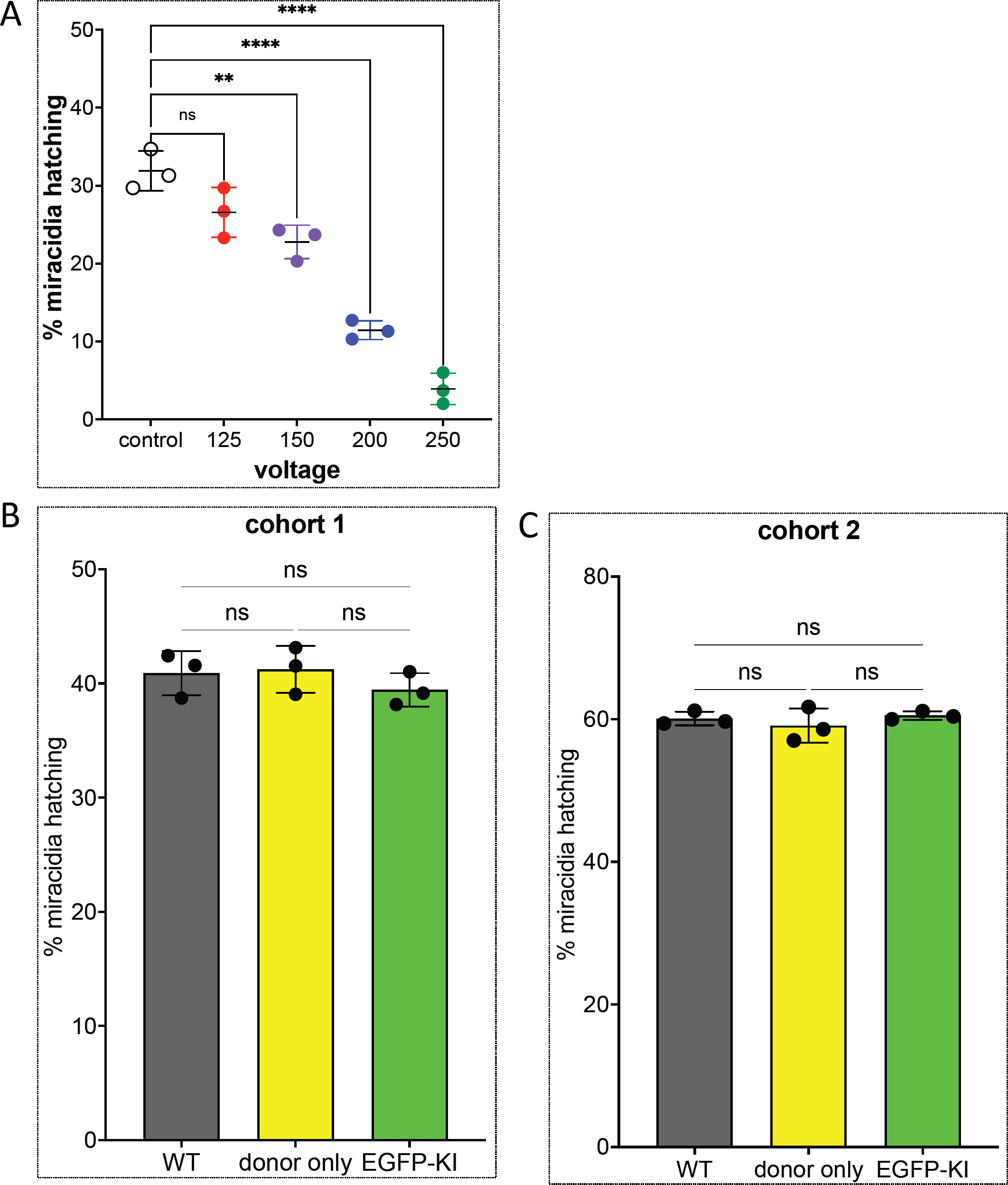
Investigation of miracidial hatching. Panel A, percent miracidia hatched following transfection by electroporation of multiple RNPS at each of 125, 150, 200 and 250 volts Survival and/or larval growth inside the egg in the 125 volts group was not significantly affected in comparison to the control group; 26.6±3.2%, 125 V group, 31.9±2.6%, control, non-electroporated group. By contrast, hatching progressively decreased as voltage increased: 150 V, 22.8±2.2%; 200 V, 11.4±1.2%; 250 V, 3.9±2.0%, respectively (*P* ≤0.001, two-way ANOVA). Panels B and C, differences were not seen in hatching rate among the treatments group of the control (WT), donor only and EGFP KI treatment groups. The experiment was performed twice with a different cohort of schistosome eggs each time, here termed cohort 1 (B) and cohort 2 (C). In both B and C, each treatment included two technical replicates, each replicate of ∼1,000 eggs. Hatching rate was established based on three blinded counts for each replicate.

**Table S1.**
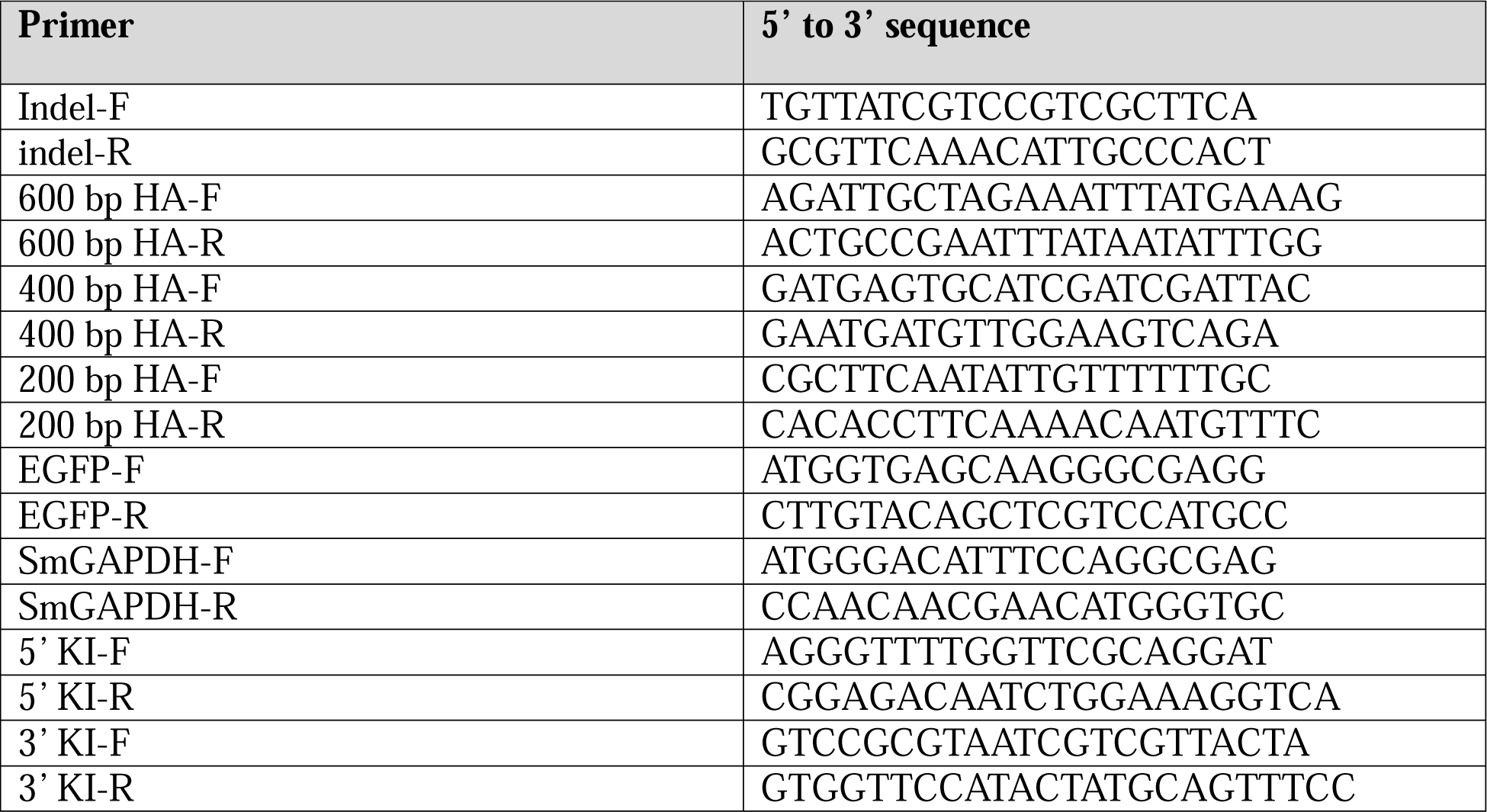
Names and nucleotide sequences of primers used in PCRs to investigate programmed knockout and knock-in.

